# Consensus Finder web tool to predict stabilizing substitutions in proteins

**DOI:** 10.1101/2020.06.29.178418

**Authors:** Bryan J. Jones, Chi Nok Enoch Kan, Christine Luo, Romas J. Kazlauskas

## Abstract

The consensus sequence approach to predicting stabilizing substitutions in proteins rests on the notion that conserved amino acids are more likely to contribute to the stability of a protein fold than non-conserved amino acids. To implement a prediction for a target protein sequence, one finds homologous sequences and aligns them in a multiple sequence alignment. The sequence of the most frequently occurring amino acid at each position is the consensus sequence. Replacement of a rarely occurring amino acid in the target with a frequently occurring amino acid is predicted to be stabilizing. Consensus Finder is an open-source web tool that automates this prediction. This chapter reviews the rationale for the consensus sequence approach and explains the options for fine-tuning this approach using *Staphylococcus* nuclease A as an example.

## Introduction

Protein stabilization is a common protein engineering goal (Kazlauskas, 2018). More stable proteins enable their use as biocatalysts in stressful environments like high temperatures, solutions containing organic cosolvents, or high substrate or product concentrations. Increased stability also extends the useful lifetime of proteins at normal temperatures and their stability to storage. More stable proteins often avoid forming inclusion bodies when overexpressed in bacteria and therefore are easier to manufacture. More stable proteins can tolerate destabilizing substitutions that confer beneficial traits like altered substrate specificity and are a better starting point for protein engineering.

The consensus sequence approach (Porebski & Buckle, 2016; Lehmann, Pasamontes, Lassen, & Wyss, 2000; Lehmann et al., 2002) is one of the easiest and most reliable approaches to stabilizing proteins. It is easy because it requires only the protein sequence, while most other methods also require a protein structure. Typically about half of the substitutions predicted to be stabilizing by the consensus approach are correct (see Table 1 below), which is higher than most computational methods. For example, only ~10% of the substitutions predicted to be stabilizing by Rosetta and FoldX were correct (Floor et al., 2014).

**Table 1.** Selected examples of single consensus substitutions to stabilize proteins.

| protein | how consensus sequence defined | success rate<sup>a</sup> | reference |
| --- | --- | --- | --- |
| VK domain of antibody McPC603 from mouse | ~200,000 VK sequences, highly similar | 6/10 | Steipe, Schiller, Plückthun, & Steinbacher, 1994 |
| human tumor suppressor protein p53 | 22 homologs | 2/17/1 | Nikolova, Henckel, Lane, & Fersht, 1998 |
| GroEL chaperone from *Escherichia coli* | 130 homologs | 13/5/16<sup>b</sup> | Wang, Buckle, Foster, Johnson, & Fersht, 1999 |
| SH3 domain from yeast actin protein 1 | 266 homologs, 27% average identity | 3/8 | Rath & Davidson, 2000 |
| penicillin G acylase from *Escherichia coli* | 8 homologs; 29-88% identity<sup>c</sup> | 10/20 | Polizzi, Chaparro-Riggers, Vazquez-Figueroa, & Bommarius, 2006 |
| isopropylmalate dehydrogenase from *Thermus thermophilus* | 18 homologs | 4/8 | Watanabe, Ohkuri, Yokobori, & Yamagishi, 2006 |
| glucose 1-dehydrogenase from *Bacillus subtilis* | 12 homologs; 25-61% identity<sup>c</sup> | 11/24 | Vázquez-Figueroa, Chaparro-Riggers, & Bommarius, 2007 |
| triosephosphate isomerase from *Saccharomyces cerevisiae* | 781 homologs | 12/23 | Sullivan et al., 2012 |
| lipase T6 from *Geobacillus stearothermophilus* | 9 homologs; 35-56% identity<sup>d</sup> | not available | Dror, Shemesh, Dayan, & Fishman, 2014 |
| esterase from tobacco | 116 homologs, <90% identity | 8/12 | Jones, Lim, Huang, & Kazlauskas, 2017 |
| amine transaminase from *Aspergillus terreus* | 90 homologs | 15/17 | Xie et al., 2019 |

<sup>a</sup> stabilizing/destabilizing or stabilizing/neutral/destabilizing
<sup>b</sup> The residues for replacement were those where <35% of the homologs contain the target residues. Decreasing the threshold to 20% increased the success rate to 13 stabilizing, 2 neutral, 3 destabilizing.
<sup>c</sup> exclude locations near active site; exclude substitutions thought to disrupt secondary structure, H-bonds or salt bridges
<sup>d</sup> exclude locations in active site and lipase lid

The consensus sequence approach starts by searching a database for the sequences homologous to the target sequence. These sequences are aligned with the target and the most frequently occurring amino acid at each position is defined as the consensus sequence (Fig. 1). Potentially-stabilizing substitutions are those that replace a rarely occurring amino acid in the target protein with a frequently occurring amino acid from the consensus sequence.

**Fig. 1.**
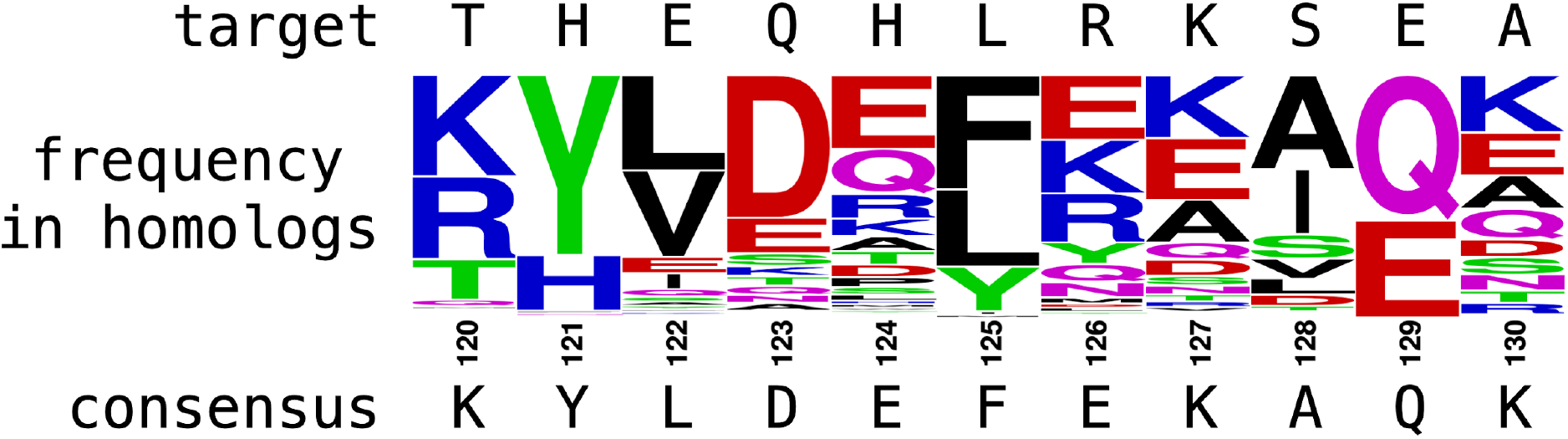
The most frequently occurring amino acid in a multiple sequence alignment defines a consensus sequence. The target sequence (*Staphylococcus* nuclease A) was aligned with 351 homologs. The height of the letters in the logo plot reflects the frequency of the amino acid in the multiple sequence alignment. The sequence of the most common amino acid at each position is the consensus sequence. This consensus sequence predicts that an H124E substitution would stabilize this protein since 30% proteins in the alignment have E, but only 1% have H. Unfolding experiments showed that this substitution slightly stabilized *Staphylococcus* nuclease (ΔΔ*G*_*unfolding*_ = +0.2 kcal/mol; Schwehm, Fitch, Dang, García-Moreno, & Stites, 2003). The web tool at weblogo.berkeley.edu generated the logo plot of amino acid frequencies.

Many groups used the consensus sequence approach to find single amino acid substitutions that stabilize their target protein, Table 1. Each application varied in the details. For example, some groups used a handful of homologous sequences (8-12) to define the consensus sequence, while others used hundreds (781) or even thousands (~200,000) of sequences. Some groups used only sequence information, while others included structural information. For example, one group excluded substitutions near the active site and those thought to disrupt the secondary structure, H-bonds, or salt bridges (Polizzi, Chaparro-Riggers, Vazquez-Figueroa, & Bommarius, 2006; Vázquez-Figueroa, Chaparro-Riggers, & Bommarius, 2007). About half of the consensus replacements prove to be stabilizing, but some of these increases were small. For example, only two of the 10 stabilizing substitutions identified for penicillin G acylase increased the half-life by more than a factor of three (Polizzi, Chaparro-Riggers, Vazquez-Figueroa, & Bommarius, 2006).

When stabilizing substitutions act independently, then combining them yields an additive increase in the stabilization. Five different combinations of four stabilizing substitutions in tumor suppressor protein p53 yielded near additive stabilization (Nikolova, Henckel, Lane, & Fersht, 1998). Two different combinations of six stabilizing substitutions in a chaperone protein yielded approximately additive stabilization (Wang, Buckle, Foster, Johnson, & Fersht, 1999). In contrast, combining fourteen stabilizing substitutions in triosephosphate isomerase destabilized the protein compared to wild type indicating non-additive behavior (Sullivan et al., 2012). Substitutions far apart from each other are more likely to act independently than nearby substitutions.

Several web tools automate the prediction of stabilizing substitutions using the consensus sequence approach: Consensus Finder (Jones, Lim, Huang, & Kazlauskas, 2017), FireProt (Musil et al., 2017), and PROSS (Goldenzweig et al., 2016). This paper focuses on Consensus Finder using nuclease A from *Staphylococcus aureus* as a test example. It is a small (149 amino acids), single-domain protein and its x-ray structure has been solved (pdb id = 1ey0). The active site residues involved in catalysis are Arg35, Glu43, and Arg87 together with a calcium ion bound to Glu21, Glu40, and Thr41. Researchers have measured the stabilizing effect of many substitutions for this protein (Byrne & Stites, 2007), but many others remain unmeasured so most of the substitutions predicted by Consensus Finder have not been checked by experiments.

## Rationale

The rationale for the consensus sequence approach is that evolution conserves amino acids that contribute to fitness and one aspect of fitness is protein stability. Conserved amino acid residues are hypothesized to be more likely to stabilize the protein fold than non-conserved residues.

One can express this notion quantitatively by extending the idea of a Boltzmann distribution to protein evolution, in particular to a sequence alignment of proteins. The Boltzmann distribution, developed to describe gas particles at equilibrium, is a probability (or frequency) distribution that varies exponentially with the energy of the state. States with lower energy are more likely to be occupied. The probability of a gas particle, *p*_*i*_, to have energy *ε*_*i*_ decreases exponentially as *ε*_*i*_ increases, eq. 1, where *k* is Boltzmann’s constant, and *T* is temperature.

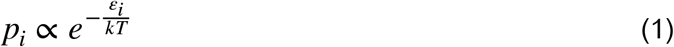

Rearrangement of eq 1 yields eq 2, which predicts the energy of the state from its frequency of occurrence.

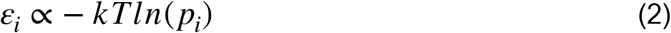

One extension of the Boltzmann distribution can accurately predict protein structures. Statistical potentials are Gibbs-energy functions derived from the frequency of occurrence of conformations in a database of structures (Shortle, 2003). If the collection of conformations in the database is a random collection at equilibrium, then more stable conformations will occur more frequently than less stable conformations. The Rosetta force field for predicting protein structures is a statistical potential because it ranks conformations according to how frequently they occur in the Protein Data Bank, eq 3.

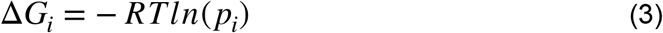

Here the gas constant, *R*, replaces Boltzmann’s constant, *k*, to calculate energies per mole instead of per particle.

The consensus sequence approach for protein stabilization further extends Boltzmann’s energy distribution beyond protein conformations to amino acid frequencies. The homologous protein sequences chosen for comparison are assumed to be equivalent to a random collection of proteins with this fold at equilibrium. At each position, amino acids that contribute more to the stability of the fold will occur more frequently than those that contribute less to stability.

To predict the stabilizing effect of a target to consensus substitution, ΔΔ*G*_*unfolding,target*→*consensus*_, one subtracts the predicted stabilizing effect of the existing amino acid from the predicted stabilizing effect of the consensus amino acid.

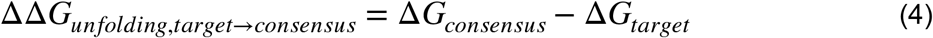

Combining equations 3 and 4 yields eq 5 where the predicted stabilization varies with the natural logarithm of the ratio of abundances of the proposed replacement amino acid from the consensus sequence and the abundance of the existing amino acid in the target.

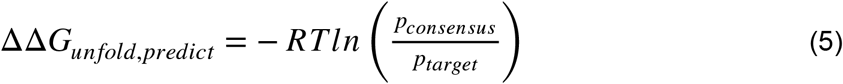

For example, if the consensus amino acid occurs in 70% of the sequences in the multiple sequence alignment, while the existing amino acid occurs in only 10% of the sequences in the multiple sequence alignment, then the predicted stabilization effect of the replacement is 4.9 kJ/ mol (1.2 kcal/mol). Since changes in the free energy of unfolding shift the equilibrium constant for unfolding, eq 6, this is equivalent to predicting that this substitution would increase the lifetime of the protein sevenfold.

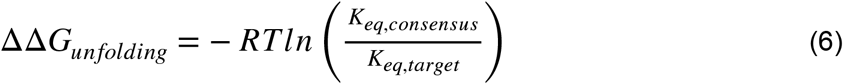

This application of the Boltzmann distribution to amino acid frequencies involves approximations that ignore the known characteristics of proteins. First, one assumes that the collection of homologous sequences reflects a random distribution. Proteins do not appear randomly but evolve from existing proteins, so collections of homologs usually contain clusters of similar sequences. Consensus Finder includes calculations to minimize this phylogenetic bias and create a more even distribution of homologous sequences. Second, evolution does not select amino acids only for stability, but also for specific functions like catalysis. One way to minimize the identification of functionally conserved amino acids as stabilizing is to restrict the homologs to those with the same function. For example, the homologs of penicillin acylase in Table 1 were all penicillin acylases (Polizzi, Chaparro-Riggers, Vazquez-Figueroa, & Bommarius, 2006). This restriction limits the comparison to well-characterized homologs. Consensus Finder includes all homologous sequences in the comparison. Third, one assumes that each amino acid contributes to stability independently, but some amino acids contribute to stability not independently but by cooperative interactions like the formation of ion pairs. Replacement of only one residue in a pair of covarying residues creates an unfavorable interaction, so the stability will be lower than predicted from the frequency of occurrence. Excluding co-varying amino acid residues improved prediction of stabilizing residues in triosephosphate isomerase (Sullivan et al., 2012), but Consensus Finder does not exclude covarying residues. Finally, the derivation above assumes that all amino acids occur with equal frequency, but amino acids occur in proteins with unequal frequencies (Lobry & Gautier, 1994). For example, leucine occurs more frequently than tryptophan. Consensus Finder does not adjust the prediction of stabilizing substitutions with this natural frequency since the merit of such an adjustment is unknown.

Several groups have tested the relationship in eq. 6 experimentally and found a moderate correlation between mutant stability and the degree of conservation of the introduced amino acid. Substitutions within β-turns in the variable domain of an immunoglobulin correlated moderately with stability: r = 0.65; eight measurements (Ohage, Graml, Walter, Steinbacher, & Steipe, 1997). Substitutions in SH3 domains correlated better with stability (r = 0.78, 26 measurements, Di Nardo, Larson, & Davidson, 2003), but this correlation excluded nine outliers. Six of these outliers involved covarying residues and three involved functional residues. Neither group adjusted for the differing natural frequencies of amino acids.

Proteins in Nature need to be only stable enough to fulfill their biological function; proteins with stabilities above a certain threshold will have no further selection advantage. Since random substitutions are more likely to be neutral or destabilizing, proteins will lose stabilizing substitutions beyond those required for adequate stability. Each homolog in the protein family has adequate stability but may acquire this stability with different subsets of stabilizing substitutions. By identifying all the stabilizing contributions by statistical analysis, one can engineer a protein that goes beyond adequate stability toward maximal stability.

## Required input: the amino acid sequence of the target protein

The only required input for Consensus Finder is the protein sequence of the target protein. This sequence should be in a FASTA format, which is a plain text file, often named using .txt or .fasta as the extension, but this extension is not required. A plain text contains only text without formatting information (font type, bold, color, etc.). The FASTA format means that the text file should contain at least two lines. The first line starts with ‘>’ (greater than symbol) and contains a description of the sequence. This is followed by one or more lines containing single-letter codes for the protein sequence. This file is uploaded to Consensus Finder.

The source of the FASTA-formatted plain text file for the target protein is usually a protein sequence database. For example, Figure 2A shows the FASTA file for *Staphylococcus aureus* nuclease A retrieved from the UniProtKB database (https://www.uniprot.org/uniprot/).

**Figure 2.**
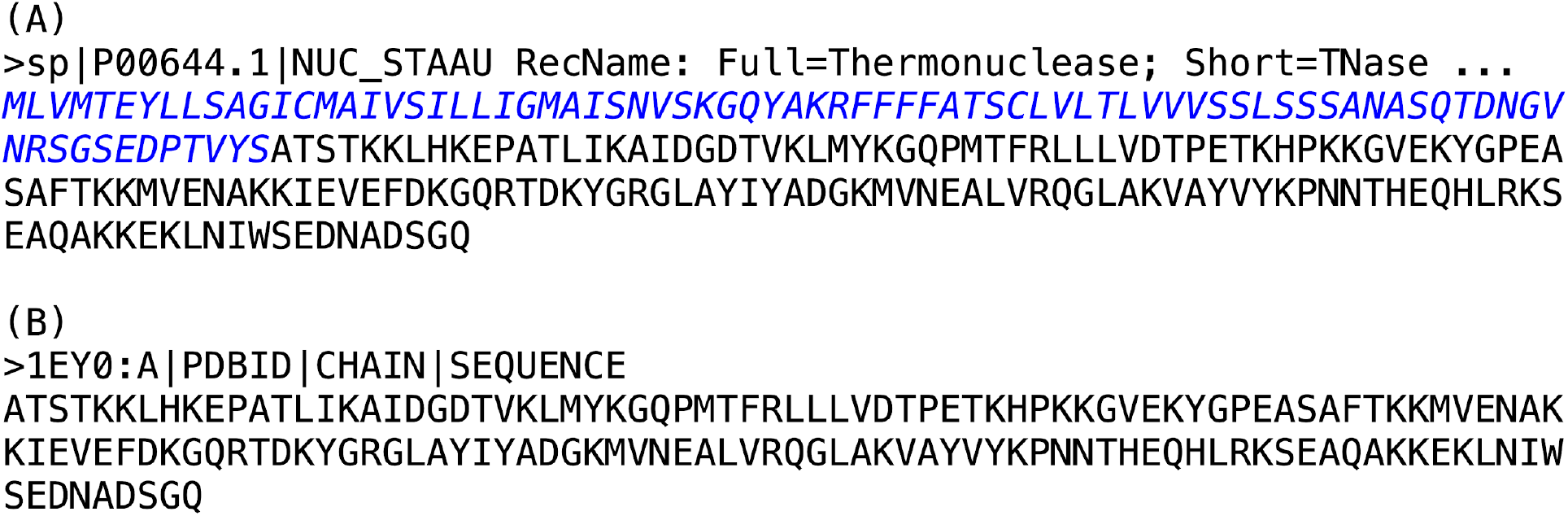
Two FASTA-formatted protein sequences for *Staphylococcus* nuclease A. The FASTA format consists of at least two lines. The first line starts with > and describes the contents. The following lines are the protein sequence. This sequence may be broken into multiple lines or all on a single line. (A) The protein sequence retrieved from the UniProtKB database includes the entire translated sequence including the signal sequence and propeptide (blue text color and italics added for clarity; there is no text formatting in the FASTA file). (B) The sequence retrieved from the Protein Data Bank contains only the mature protein sequence. The mature protein sequence is preferred for predicting stabilizing substitutions.

Protein sequences may differ between databases. For *Staphylococcus* nuclease A, the protein sequences differ between the UniProtKB and the Protein Data Bank. The sequence from UniProtKB contains all 231 translated amino acids including the initial 82 amino acids corresponding to the signal sequence and the propeptide. The sequence from the Protein Data Bank contains only 149 amino acids corresponding to the mature protein. It matches residues 83-231 from the UniProtKB sequence. Since the goal is the stabilization of the mature protein, the appropriate sequence for this example is the mature protein sequence found in the Protein Data Bank.

## Prediction with default settings

Using Consensus Finder with the default setting only requires specifying the protein sequence. One way to specify this sequence is to click on ‘Browse…’ and choose a file from your computer, Figure 3. This example uses a FASTA-formatted file, which was previously downloaded from the Protein Data Bank. Figure 2B above shows the contents of this file. Alternatively, if the Protein Data Bank contains a structure of the target protein, then the pdb id of the structure can be entered instead. In this case, Consensus Finder will automatically download the FASTA-formatted protein sequence associated with the structure. Consensus Finder includes a copy of the input target protein sequence in the complete output allowing the user to check the sequence used for the calculations.

**Fig. 3.**
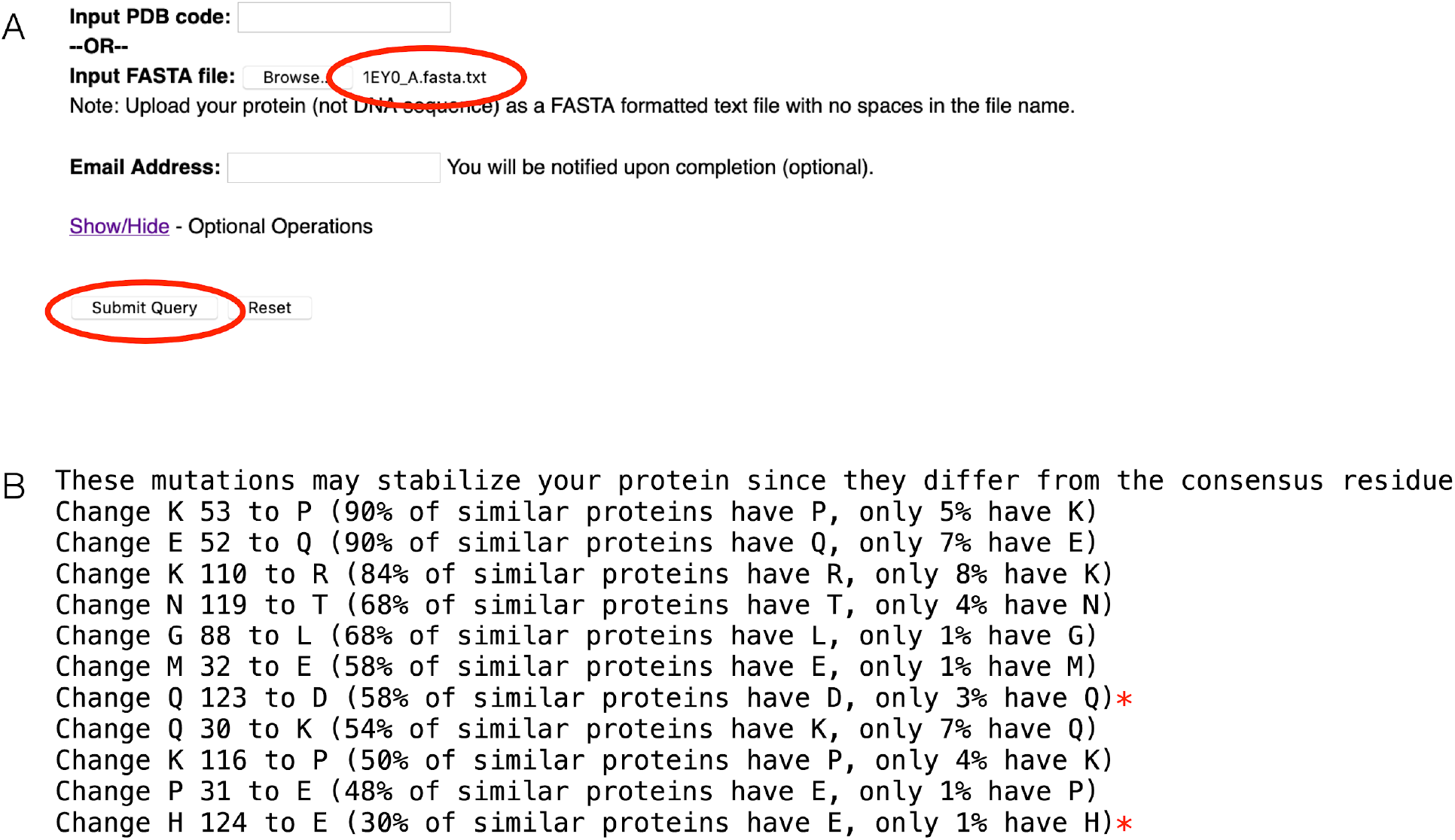
Predicting stabilizing substitutions using Consensus Finder and default settings. (A) The input screen shows that the user has selected a text file (1EY0_A.fasta.txt) on their computer for upload. Figure 2B above displays the contents of this file. (B) The output screen is a list of eleven potentially-stabilizing substitutions. Experimental data are available only for the two substitutions marked with an asterisk and suggests that they are both stabilizing as predicted. Downloading the complete output will include a file named mutations.txt that contains this list of predictions.

To generate this prediction Consensus Finder searched for homologs and defined a consensus sequence based on 362 homologs. These are not displayed, but are available in the complete output as files named alignment.fst and consensus.fst, respectively. Consensus Finder displays a suggestion of eleven possibly-stabilizing substitutions ranked according to the frequency of the replacement amino acid within the multiple sequence alignment. The first suggestion is K53P since 90% of the proteins in the sequence alignment have proline at position 53, but the target protein has a lysine. Introducing proline often stabilizes proteins because its ring structure limits the flexibility of the unfolded protein (Watanabe & Suzuki, 1998). The predicted substitutions suggest introducing proline at two positions, 53 and 116. It also suggests removing proline at position 31. The substitution might be stabilizing if proline fits poorly in the folded protein at this position. The suggested replacement of glycine at position 88 also expected to stabilize the protein because the replacement amino acid will limit flexibility of the unfolded protein as compared to the highly flexible glycine (Matthews, Nicholson, & Becktel, 1987). Experimental data are available for only two of the eleven predicted substitutions. The H124E variant is 0.2 kcal/mol more stable than wild type to denaturation with guanidinium chloride (Schwehm, Fitch, Dang, García-Moreno, & Stites, 2003). The stability of the Q123D variant was not measured, but the x-ray crystal structure showed a slightly higher resolution and ionic interactions suggesting that it may be more stable (Jeliazkov, Robinson, García-Moreno, Berger, & Gray, 2019).

Including the propeptide and signal sequence in the target changes the consensus sequence, but predicts the same eleven substitutions. The search for homologs finds additional sequences that match the propeptide and signal region, so the alignment includes 36 more sequences than the alignment of the homologs of the mature protein (398 instead of 362). The consensus sequence is longer because it includes the propeptide and signal sequence. The consensus sequence for the full sequence also differs at sixteen locations in the mature region from the consensus sequence for the mature protein. Within the first twelve amino acids, nine amino acids differ between the two consensus sequences. The adjacent propeptide led to the inclusion of homologous sequences that matched the propeptide region. Seven additional differing amino acids are scattered in the rest of the consensus sequences. Despite these differences, the predicted stabilizing substitutions are the same eleven substitutions in Fig 3B for both target sequences. For the full sequence, Consensus Finder also predicts three substitutions in the propeptide region, but these can be ignored because they are removed in the mature protein.

## Adjusting the default settings

To fine-tune the predictions, one can adjust the default settings. This section explains each calculation step that Consensus Finder uses to make the predictions, what the default settings are, and how adjusting these settings affects the prediction.

Consensus Finder first defines the consensus sequence for the target protein and next compares the consensus sequence and target protein to predict stabilizing substitutions. The steps involved in defining the consensus sequence are:

- retrieve sequences similar to the target protein from the NCBI non-redundant protein database (BLAST search)
- remove overrepresented sequences from the collection using CD-HIT
- align the sequences using Clustal Omega and calculate the frequencies of the amino acids occurring at each position
- list the most commonly occurring amino acid at each position to define the consensus sequence

The steps involved in comparing the consensus sequence and target protein to predict stabilizing substitutions are:

- identify locations that differ between the target and consensus sequences
- Is the current amino acid in the target protein rare enough and the consensus amino acid common enough that the substitution would likely be stabilizing?
- (optional: exclude amino acids close to the active site)
- list potentially stabilizing substitutions

### Defining a consensus sequence

The consensus sequence for a target protein is not unique but varies with the sequences that are included in the comparison. The first group of options, Table 2, adjusts the search for homologs and the multiple sequence alignment that defines the consensus sequence.

**Table 2.**
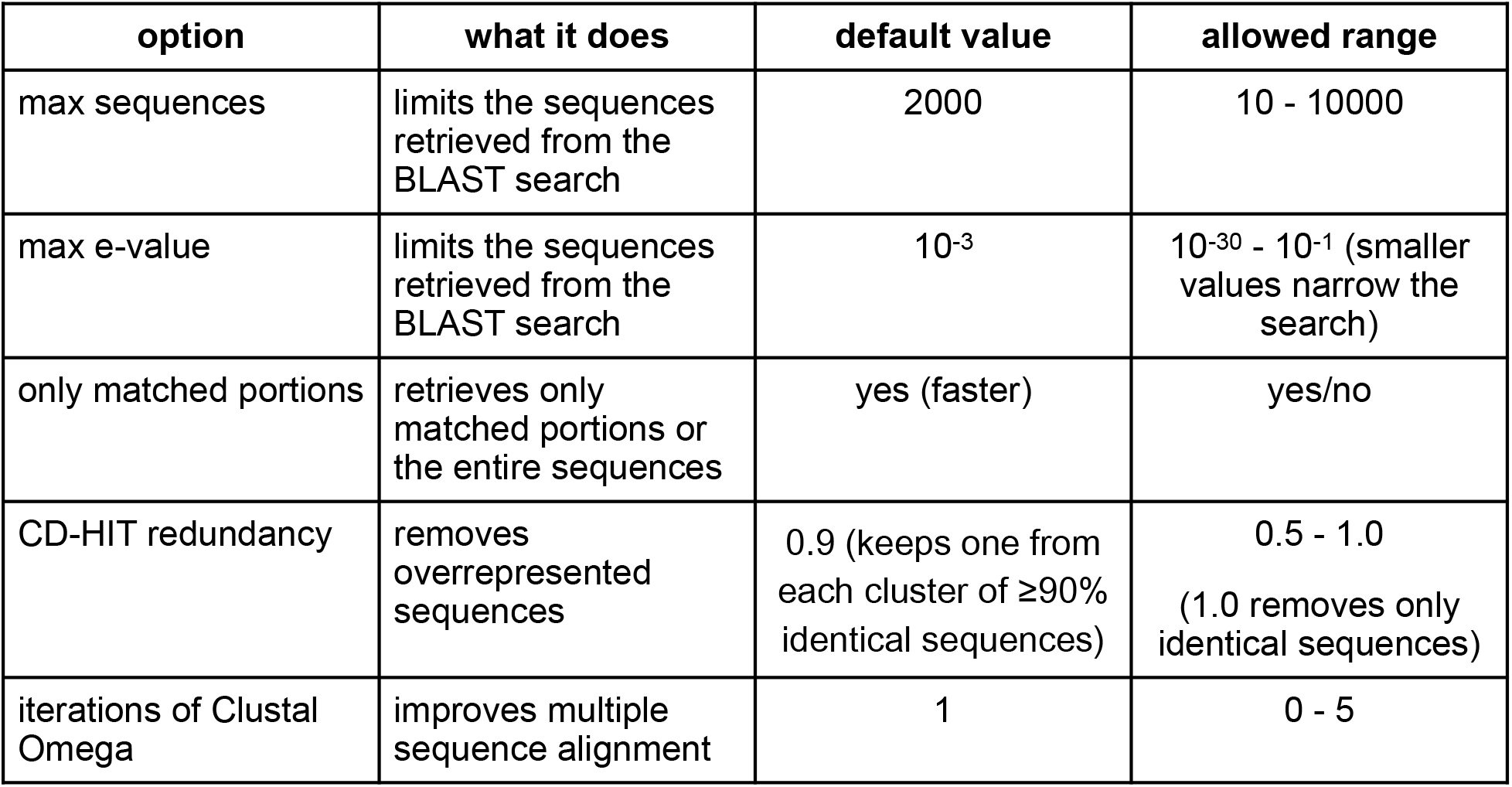
Optional settings in Consensus Finder that may alter the consensus sequence.

| option | what it does | default value | allowed range |
| --- | --- | --- | --- |
| max sequences | limits the sequences retrieved from the BLAST search | 2000 | 10 - 10000 |
| max e-value | limits the sequences retrieved from the BLAST search | $10^{-3}$ | $10^{-30}$ - $10^{-1}$ (smaller values narrow the search) |
| only matched portions | retrieves only matched portions or the entire sequences | yes (faster) | yes/no |
| CD-HIT redundancy | removes overrepresented sequences | 0.9 (keeps one from each cluster of $\geq 90\%$ identical sequences) | 0.5 - 1.0<br>(1.0 removes only identical sequences) |
| iterations of Clustal Omega | improves multiple sequence alignment | 1 | 0 - 5 |

**Table 3.** Options in Consensus Finder for setting the threshold for predicting a stabilizing substitution.

| option | what it does | default value | allowed range |
| --- | --- | --- | --- |
| ratio | sets threshold for suggesting a stabilizing substitution | 7 | 1 - 100 (1 allows substitution with equally occurring or more common amino acids) |
| conservation threshold | an alternative measure to set a threshold | blank (use only ratio) | 0.05 - 0.99 |

**Table 4.** Required and optional settings to exclude substitutions near the active site.

| setting | what it does | default value | allowed range |
| --- | --- | --- | --- |
| pdb id (required) | identifies structure of target protein | no default | valid pdb id, e.g., 1ey0 |
| active site residue (required) | defines center of active site | no default | 1 - last amino acid residue in chain |
| pdb chain (optional) | choose chain that defines the active site | A (first chain) | A, B, C, up to the last chain in the structure |
| size of active site (optional) | defines radius of active site | 5 Å | 2 - 20 Å |

Consensus Finder starts by retrieving sequences similar to the target sequence using a BLAST search of the NCBI non-redundant protein database. The default settings of Consensus Finder enable a relatively broad search to extend to distantly related sequences (max e-value = 10^−3^) and return a maximum of 2000 of the most closely related sequences. Either the max e-value setting or the maximum number of sequences will limit the number of sequences retrieved. If the database contains many related sequences, then the maximum number will be reached quickly to stop the retrieval. The resulting collection will be closely related to the target. If the database contains fewer related sequences, then the BLAST search includes more distantly related sequences and will be limited by the max e-value. Increasing the maximum number of sequences and the maximum e-value expands the search to include more distant sequences.

The optimum range of sequences to include is currently unknown. The comparison sequences should include enough variation to distinguish variable sites from conserved sites. If the target sequence is 200 aa acids long, then 2000 examples of substitutions correspond to an average of 10 substitutions at each position. To get these 2000 examples, one could retrieve 200 sequences each containing an average of 10 substitutions (or 95% identical to the target) or fewer sequences containing more substitutions. For the examples in Table 1, researchers tended to use a large number of sequences when they were very similar to the target, but fewer sequences when they were less similar. For example, defining the consensus sequence for an antibody domain (110 aa) used ~200,000 highly similar sequences, while defining the consensus for an SH3 domain (60 aa) used only 266 sequences, but they differed significantly from the target sequence (average of 27% identity).

Sequences too distant from the target sequence may lower the reliability of predictions. Although many amino acid residues contribute to stability independently, some act cooperatively. For example, the formation of an ion pair requires two partners. Exchanging an amino acid residue between close homologs maintains most of the same pairwise interactions in both homologs, but exchanging an amino acid residue between distant homologs creates many new pairwise interactions (Govindarajan et al., 2003; Lunzer, Golding, & Dean, 2010). Some of these interactions may be deleterious and therefore lower the reliability of prediction when too distant homologs are included.

The default option to ‘Use only matched portions, not complete sequences’ limits the retrieval of sequences to only those portions that match the target sequence. The alternative, to also download the non-matching portion of the sequences, slows down the calculation. The non-matching portions may form different structures and are unlikely to improve the prediction.

Next, Consensus Finder uses CD-HIT (Li, Jaroszewski, & Godzik, 2001; Fu, Niu, Zhu, Wu, & Li, 2012) to remove overrepresented sequences. These might be sequences from more intensely studied organisms. CD-HIT groups similar sequences into clusters and then chooses a random sequence from each cluster as a representative of that cluster. The default value of 0.9 creates clusters of sequences where 90% of the amino acids are identical. This pruning yields a collection of sequences that are less than 90% identical. Increasing the default value reduces the amount of pruning up to a maximum of 1.0, which removes only identical sequences. Setting the redundancy value high (small clusters) risks retaining bias toward overrepresented subgroups while lowering the redundancy value (large clusters) discards increasing numbers of sequences and the variation information they contain. For *Staphyloccocus* nuclease, a maximum sequence setting of 1000 sequences in the BLAST search yielded 65 sequences in the final multiple sequence alignment, while a maximum sequence setting of 2000 sequences (default value) yielded 352 sequences in the final alignment, Fig 4. The final number of sequences was smaller than the maximum value because CD-HIT removed sequences that were ≥90% identical to each other. The resulting consensus sequence differs at five positions among the eleven positions in this region. Two of the stabilizing substitutions predicted using the default values (Q123D, H124E, see Fig 3 above) occur in this region. The prediction using the smaller set of sequences also predicts the Q123D substitution to be stabilizing but differs at position 124. The smaller set of sequences predicts that the H124K substitution would be stabilizing. This substitution has not been tested.

**Figure 4.**
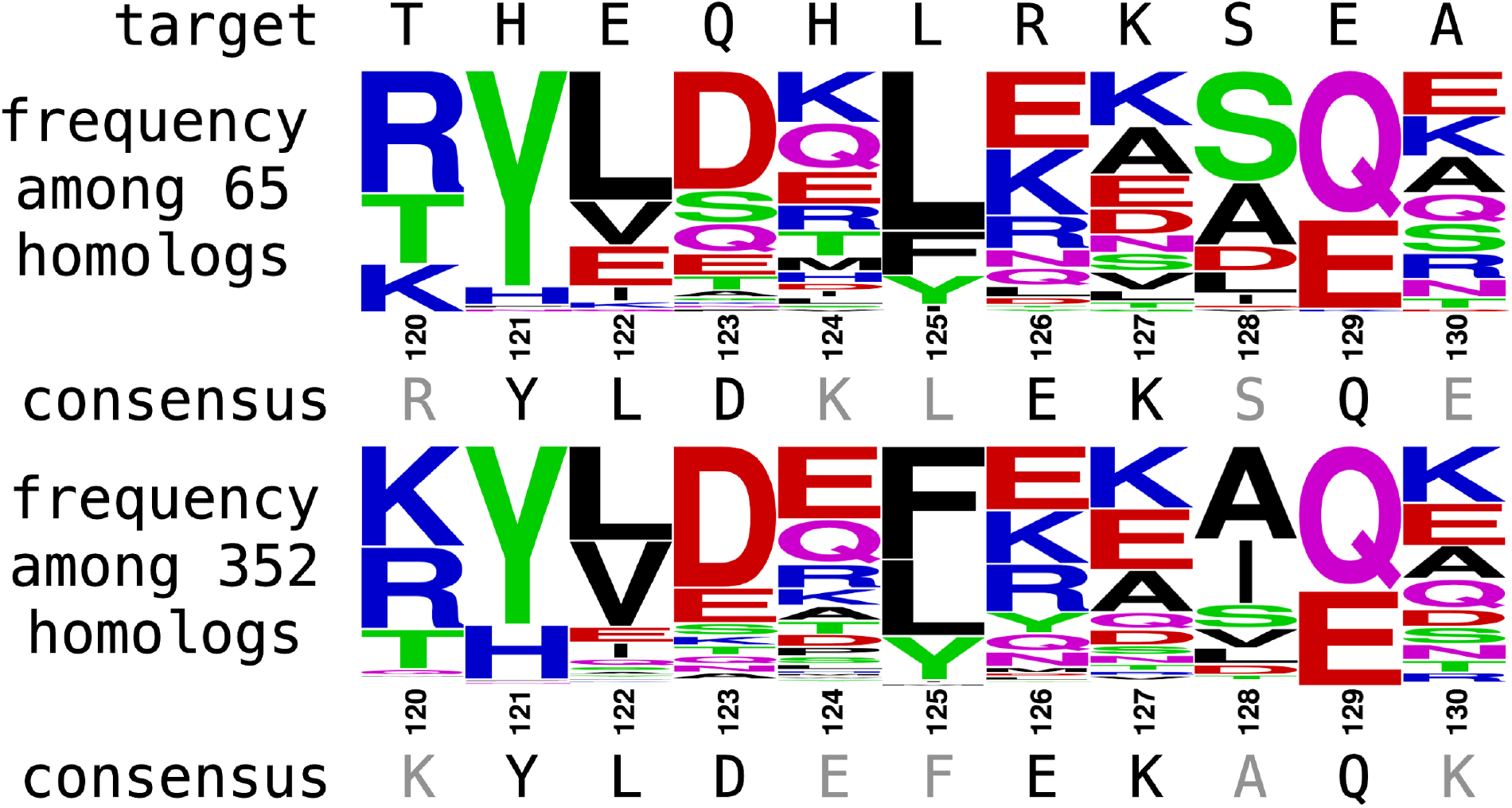
Different sets of homologs yield different consensus sequences. The consensus sequence for *Staphylococcus* nuclease A generated from 65 homologs (top) differs from the one generated from 352 homologs (bottom) at five of the eleven positions (gray text). Upon increasing the number of homologs, the most frequent amino acid switched from R to K at position 120, from K to E at position 124, from L to F at position 125, from S to A at position 128, and from E to K at position 130. The different consensus sequences predict different stabilizing substitutions for *Staphylococcus* nuclease in this region.

Clustal Omega (Sievers & Higgins, 2018) aligns the sequences into a multiple sequence alignment. Iterations improve the multiple sequence alignment but add calculation time. Multiple sequence alignments may vary depending on the early sequences chosen for the alignment. Iterations of alignment use the previous alignment as a better starting point for a new alignment. The default value is one iteration, meaning the alignment is refined once. The option is to refine this alignment up to five times.

In the case of *Staphyloccocus* nuclease, changing the number of iterations changes the resulting consensus sequence slightly, Fig 5. Increasing from no iterations to one iteration changes the location of a two-amino-acid gap from after position 10 for no iterations, to before position 1 with one iteration. Thus, the two consensus sequences differ at four locations. Further increasing the number of iterations from one to five changes the consensus amino acid at position 80. It is proline for zero and one iteration, but changes to glutamate with five iterations.

**Figure 5.**
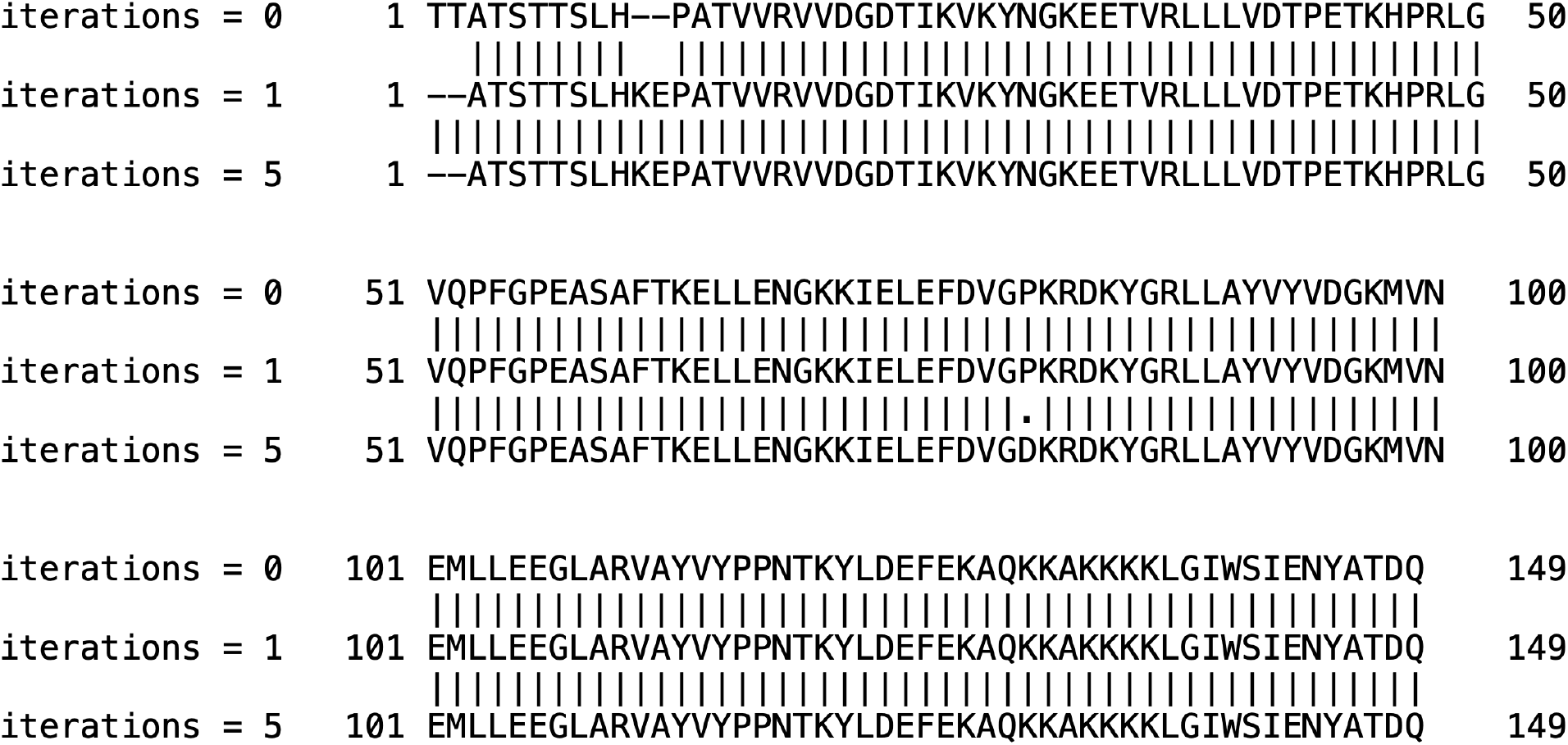
Increasing the number of iterations of the multiple sequence alignment for *Staphylococcus* nuclease slightly changes the resulting consensus sequence. Iterations improve sequence alignments by using the previous multiple sequence alignment as the starting point for a new alignment.

Finally, Consensus Finder trims the multiple sequence alignment to include only positions in the target sequence. Some of the retrieved sequences contain insertions compared to the target sequence, so the resulting multiple sequence alignment will be longer, often much longer, than the target sequence. The goal of the alignment is to compare each position of the target sequence to the corresponding position in the multiple sequence alignment, so insertions do not contain useful data. The trimming step shortens the multiple sequence alignment to the same length as the target sequence.

### Predicting stabilizing substitutions from the consensus sequence

Consensus Finder predicts selected substitutions that are most likely to stabilize the target protein. Consensus Finder compares the frequency with which the consensus amino acid appears in the multiple sequence alignment to the frequency with which the existing amino acid in the target sequence appears in the multiple sequence alignment. The frequency calculation groups gaps in the alignment (deletions in homologs) and non-standard amino acids (e.g., selenomethionine) as ‘other’ and omits them from the total number of amino acids at that location. The predicted change in free energy of stabilization depends on the logarithm of the ratio of these two frequencies, so the default comparison is this ratio. If the ratio is greater than seven, which corresponds to a predicted 4.9 kJ/mol (1.2 kcal/mol) increase in the free energy of unfolding (see eq 5 above), then Consensus Finder predicts that the substitution will stabilize the protein. Setting this ratio lower will suggest more substitutions while setting it higher will suggest fewer substitutions.

As an alternative, one can also set an absolute value for conservation. If the target and consensus residues differ and the consensus residue occurs with a frequency above this threshold, then Consensus Finder predicts a replacement to the consensus residue regardless of the frequency of the existing residue at a target. At low values, this approach predicts all substitutions needed to create a full consensus sequence. The rationale for the consensus sequence approach relies on the ratio of the frequencies of amino acids in target and consensus, so this absolute value approach is not recommended.

While the consensus sequence approach does not require structural information, Consensus Finder can use a structure from the Protein Data Bank to exclude substitutions near the active site, Table 5. While substitutions near the active site can stabilize a protein, they are likely to disrupt catalysis and binding, so most researchers avoid substitutions near the active site. To exclude substitutions near the active site, the user must enter the target protein sequence using the pdb id option. Second, on the Optional Operations page, the user must specify the amino acid residue corresponding to the center of the active site. Consensus Finder then excludes any amino acid that lies with 5 Å of the specified amino acid. In the case of *Staphylococcus* nuclease, excluding residues with 5 Å of R35 in the active site would exclude one substitution (G88L) from the prediction using the default setting in Fig 3 because glycine lies within 5 Å of arginine 35. One option is to expand or contract the excluded region. Consensus Finder uses chain A in the protein structure for the calculation, but the user can choose a different chain. Consensus Finder assumes that the active site lies entirely on a single chain.

## Other application of consensus sequences

The consensus sequence approach can also be used to design new proteins. Instead of making single consensus substitutions in a target protein, researchers synthesized the protein containing the consensus amino acid at every location, known as the consensus protein (Desjarlais & Berg, 1993). The goal of this full consensus approach is to create a non-natural sequence with native-like structure and function. For example, a full consensus protein design of ankyrin protein (a binding protein) yielded a stable protein with ankyrin fold, but the individual binding specificities were averaged out leaving an inert scaffold (Forrer et al., 2004).

The data from a consensus sequence analysis also identifies positions in a protein that tolerate substitutions. These variable positions are sometimes called mutational hot spots. Creating libraries of substitutions known to occur in homologs can reshape substrate binding sites for new substrates without compromising the stability of the protein (Jochens & Bornscheuer, 2010).

## Acknowledgments

This work was supported by the National Science Foundation (CBET 1930825) and the University of Minnesota.

